# Forest structural diversity determines tree growth synchrony in response to climate change

**DOI:** 10.1101/2023.03.17.532989

**Authors:** J. Astigarraga, J. Calatayud, P. Ruiz-Benito, J. Madrigal-González, J. Tijerín-Triviño, M. A. Zavala, E. Andivia, A. Herrero

## Abstract

After centuries of deforestation, many industrialised countries are experiencing an increase in forest area and biomass due to changes in land- and forest-use since the mid-20^th^ century. At the same time, the impacts of climate change on forests are aggravating, but the interplay between past land- and forest-use (i.e. land- and forest-use legacies) and climate change in forest functioning remains elusive. Here using network theory and linear mixed models, we quantified how land- and forest-use legacies modulate tree growth synchrony in response to climate change. We analysed tree growth data from European beech (*Fagus sylvatica* L.) stands with different histories of forest management at the species’ rear edge. We found that tree growth synchrony increased following heatwaves, late spring frosts, and reduced precipitation. Interestingly, the greatest tree growth synchrony occurred in recently-established forests, while stands containing large trees and heterogeneous tree sizes showed much lower growth synchrony. Our results highlight the importance of maintaining large trees and structurally heterogeneous forests to mitigate the negative effects of climate change on forest productivity, and thereby, increase forest resilience to future forest climate risks.

## INTRODUCTION

Forests have undergone major human-induced changes and today 75% of the world’s forests are altered by humans (FAO, 2020). Anthropogenic impacts on forests can alter tree cover completely through land-use changes (e.g. deforestation, afforestation or natural regrowth; Song et al. (2018)) or partially through changes in forest-use (e.g. increases in biomass after management abandonment; Jump et al. (2017)). Furthermore, climate change is causing shifts in mean temperature, precipitation and extreme climatic events (IPCC, 2021), having direct impacts on forest functioning (Allen et al., 2010). Direct anthropogenic impacts on forests, through changes in land- and forest-use, and indirect impacts, through climate change, will largely determine future forest dynamics (McDowell et al., 2020). Quantifying the interaction between these direct and indirect changes in forest functioning is both a scientific and management priority.

Many industrialised countries are experiencing an increase in forest area, density and biomass since the mid-20^th^ century due to changes in land- and forest-use such as the abandonment of agriculture and traditional forest management (Garbarino et al., 2022; Infante-Amate et al., 2022; Song et al., 2018). Land-use changes determine forest age and structure, with an expected increase in vulnerability to climate change in recently-established forests compared to long-established forests (Alfaro-Sánchez et al., 2019; Correia et al., 2021; Mausolf et al., 2018). Forest-use changes determine forest structure and, therefore, could also influence current forest ecosystems responses to climate (Marqués et al., 2022; Perring et al., 2018; Sangüesa-Barreda et al., 2015). For example, the abandonment of forest-use could lead to structural overshoot processes in which increased tree biomass in a mild climate can lead to forest decline (i.e. growth decrease, crown die-back and increased mortality) in response to more severe climatic conditions (Jump et al., 2017; Zhang et al., 2021). However, the underlying drivers by which trees may respond differently to climate as a function of past land- and forest-use (i.e. land- and forest-use legacies) have been largely unexplored.

In the current context of climate change, warming, increased aridity and heatwaves are impacting the main demographic rates of tree species (Astigarraga et al., 2020; Peng et al., 2011; van Mantgem et al., 2009), shaping forest structure (McIntyre et al., 2015; Zhou et al., 2013), composition (Esquivel-Muelbert et al., 2019; Ruiz-Benito et al., 2017) and productivity (García-Valdés et al., 2021; Zhang et al., 2018). At the same time, variations in annual precipitation and late spring frosts are also affecting tree growth and leaf damage (Castagneri et al., 2015; Sangüesa-Barreda et al., 2021; Zohner et al., 2020). Although there is a broad scientific consensus on increased climate-induced tree mortality, the same is not true for tree growth since warming can have antagonistic effects (Allen et al., 2015; Díaz-Martínez et al., 2023). For example, warming can increase tree growth, primarily through CO_2_ fertilization and removal of low temperature limitations to photosynthesis, but excess warming may reduce tree growth due to reduced water availability and increased carbon respiration costs (Adams et al., 2009; D’Orangeville et al., 2018; Peñuelas, Ciais, et al., 2017). Tree growth is a demographic rate of paramount importance that integrates many environmental constraints of tree performance, being considered an indicator of tree vitality (Dobbertin, 2005) and that can be used as an early-warning signal of tree die-off (Cailleret et al., 2019).

In the last years, synchrony in tree growth (i.e. coincident increase in tree growth over time between different tree individuals) has received great attention by researchers worldwide (del Río et al., 2021; Shestakova et al., 2016). Several studies have shown an increasing trend in tree growth synchrony associated with pervasive climatic effects from local to regional scales (Gazol et al., 2020; Latte et al., 2015). Tree growth synchrony is widely recognized as an indicator of forest vulnerability to climate change (Boden et al., 2014; Shestakova et al., 2016), since more synchronous responses might decrease population stability and persistence (Tejedor et al., 2020). Thus, increased growth synchrony to climatic stressors exacerbates the risk of widespread negative impacts on tree vitality by losing the benefits of individual-level variability in response to climate, reducing forest resistance and resilience (Clark et al., 2012). Tree growth synchrony is generally calculated as shared variation within-population tree chronologies (Tejedor et al., 2020). This could limit the conclusions drawn from the analyses of tree growth synchrony, since different tree sizes and growth forms usually respond differently to climatic stressors (Andivia et al., 2020). Therefore, assessing individual tree growth synchrony in various forest structures resulting from different land- and forest-use legacies could provide new insights in the evaluation of forest vulnerability to climate change stressors.

In the Iberian Peninsula, the long-term interactions between humans and nature have created complex socioecological systems that shape forest structure, distribution and species composition (Blondel, 2006; Scarascia-Mugnozza et al., 2000). However, since the middle of the last century, the abandonment of agricultural activities has led to an increase in forest extension and density (Poyatos et al., 2003; Vilà-Cabrera et al., 2017). Furthermore, the substitution of firewood for fossil fuels led to the abandonment of traditional forest-use, decreasing centuries-old management techniques such as coppicing and pollarding (Infante-Amate et al., 2022; Sjölund & Jump, 2013). All these anthropogenic impacts in land and forest use could interact with the increasing effects of climate change, making the fate of Iberian forests uncertain.

Here, we analysed how land- and forest-use legacies modulate tree growth synchrony in response to climate change. We hypothesised that (i) there will be an increase in tree growth synchrony regardless of land- and forest-use legacies as precipitation, heatwaves and late spring frosts are becoming an increasingly limiting factors for tree growth; and (ii) land- and forest-use legacies with greater forest structural diversity will have a lower tree growth synchrony due to greater individual-level variability in response to climate. To this end, we used the European beech (*Fagus sylvatica*, L.) as the target species, a widely-distributed species of great economic and ecological importance that is susceptible to drought and spring frosts (Archambeau et al., 2020; Cavin & Jump, 2017; Packham et al., 2012). To test these hypotheses, we quantified tree growth synchrony in stands with contrasting land- and forest-use legacies in an area with similar biotic and abiotic conditions. We mixed 12 European beech stands with four different legacies (i.e. recently-established, long-established, recently-pruned pollards and old-pruned pollards) located in the north of the Iberian Peninsula analysing *c*. 240 tree cores. Our results provide evidence of the key role of anthropogenic legacies in determining forest ecosystems responses to climate change by altering forest structural diversity.

## MATERIALS & METHODS

### Study area

The study area is a natural forest dominated by *Fagus sylvatica*, located at the southern limit of the species global distribution (Packham et al., 2012). It is in the north of the Iberian Peninsula at the easternmost part of the Cantabrian Range (42º58’N - 2º24’W; altitude: 750-1000 m a.s.l.; Fig 1). The studied forest is mainly composed of detrital sedimentary rocks and presents an oceanic climate (*Cfb*, Kottek et al., 2006), with average precipitation of 932 mm and average temperature of 13 ºC (data from easyclimate R package (Cruz-Alonso et al., 2021; Moreno & Hasenauer, 2016; Rammer et al., 2018)). Although *F. sylvatica* is the most abundant species, other tree species characteristic of European temperate forests can also be found (e.g. *Quercus robur* L., *Q. petraea* Matt., *Ilex aquifolium* L., *Sorbus aucuparia* L.).

**Figure 1.**
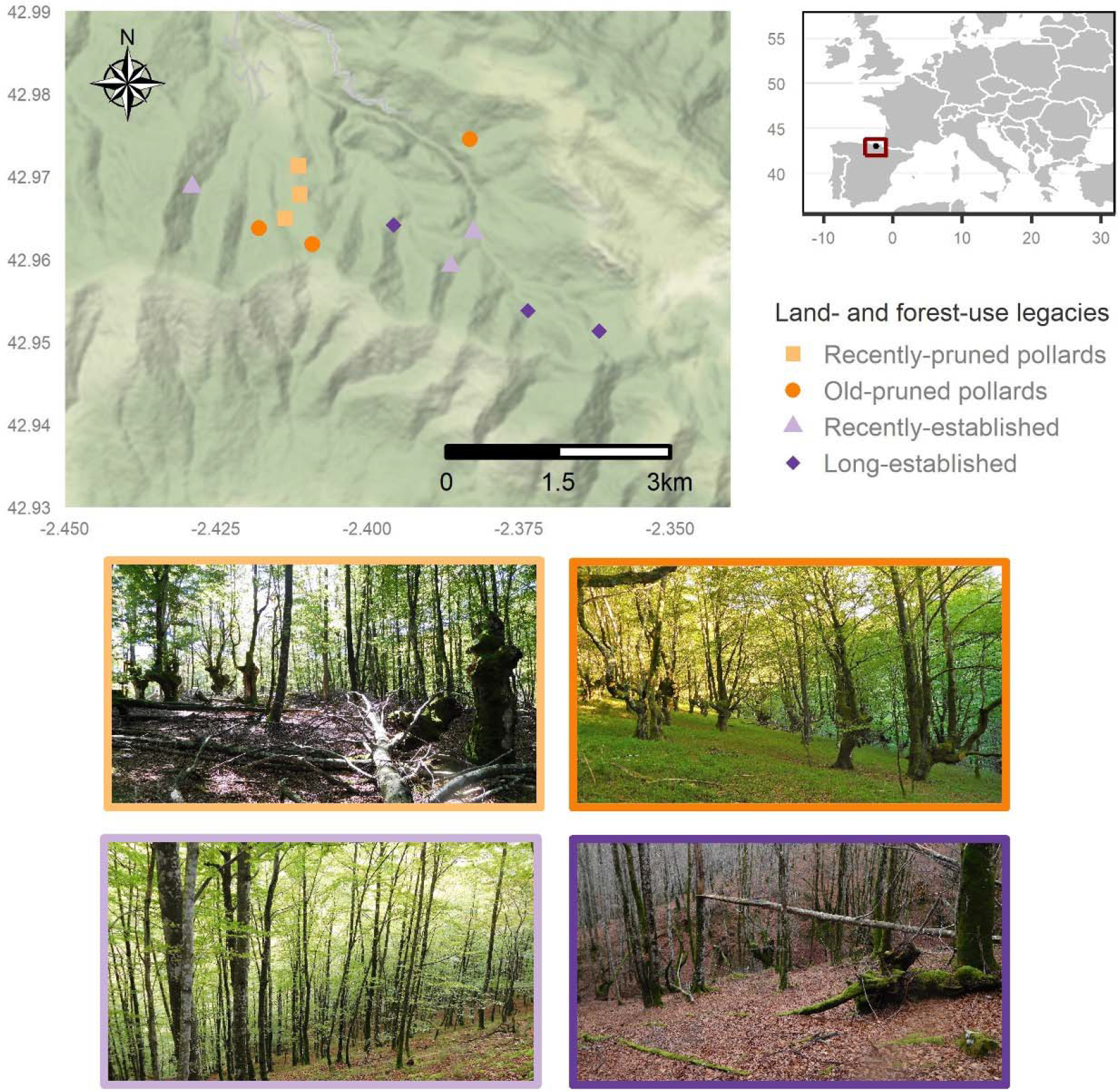
Map of the study area showing the 12 sampled stands and photographs from each legacy type. The border colors of the photographs indicate the legacy type shown in the legend.

During the last centuries, the forest was mainly used for charcoal and firewood, and its traces can be seen in the many pollards and coppiced trees that exist today in the area. Since the 1950s with the industrial transition, charcoal and firewood were replaced by fossil fuels, leading to an increase in tree density and biomass. Furthermore, agricultural and livestock activities in the region were also reduced or abandoned resulting in an increase in forest area.

### Characterisation of land- and forest-use legacies

We determined land- and forest-use legacies in the stands using aerial photographs, forest management plans and interviews to local inhabitants and forest managers. First, we distinguished recently-established from old-growth stands that were already occupied by trees in the mid-20^th^ century using aerial photographs from 1945/46 (https://www.geo.euskadi.eus/comparador-de-ortofotos/webgeo00-content/es/). Then, among stands occupied by trees prior to 1945/46, we distinguished the pollarded from high-stands. Pollarding is a traditional silvicultural technique that favours the production of a dense mass of branches above ground level, out of reach of browsing mammals, often resulting in large trees with a longer lifespan than their naturally growing counterparts (Sjölund & Jump, 2013). Among pollard stands we distinguished old-pruned pollard stands (pruned before 1970) and recently-pruned pollard stands (pruned before 1970 and in 2010 and 2014). Thus, we classified our 12 stands into (i) high-stands occupied by trees post to 1945/46 (hereafter, recently-established) and (ii) high-stands occupied by trees prior to 1945/46 (hereafter, long-established), (iii) old-pruned pollard stands, and (iv) recently-pruned pollard stands.

### Sampling

In 2020, we extracted two cores per tree in opposite directions using a Pressler increment borer and processed them using standard dendrochronology techniques (Fritts, 1976). Since each legacy type was replicated three times, we sampled 12 different stands (four legacies ×three replicates = 12 stands, Fig. 1), collecting samples from 20 individuals in each stand (four legacies ×three replicates ×20 individuals = 240 individuals). All stands were located within *c*. 1300 ha, mixing stands with different land- and forest-use legacies –although recently-pruned pollards were close to each other since the pruning was done only in that area. We georeferenced each sampled tree and measured the diameter at breast height (d.b.h.). Then, we measured ring widths using trini R package (Astigarraga et al., 2022) and averaged growths per year from both cores from each individual tree. All cores were visually cross-dated following the procedures described by Yamaguchi (1991).

### Climate data

We used daily data on precipitation and minimum and maximum temperatures from easyclimate R package (Cruz-Alonso et al., 2021; Moreno & Hasenauer, 2016; Rammer et al., 2018). We calculated total precipitation, number of late spring frosts (i.e. number of days below 0 ºC in meteorological spring –from 1^st^ of March to 31^st^ of May) and number of heatwaves (i.e. number of days above 32 ºC in meteorological summer –from 1^st^ of June to 31^st^ August) annually from 1950 to 2020.

### Statistical analyses

#### Tree growth synchrony

We used a network approach to evaluate time variations in tree growth synchrony. We built networks where nodes were trees and a link between a pair of trees was established if their growths were significantly correlated over time (*P* < 0.05, based on Spearman rank correlations. If trees of a stand were growing synchronically, tree growth networks would be densely connected, indicating that several trees showed similar growth patterns over time. Thus, network density or connectivity provides a direct measure of forest growth synchrony. We calculated network connectivity as the number of realized links relative to the number of potential links, 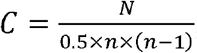, where *N* denotes the number of observed links and *n* the number of nodes. To evaluate growth synchrony over time we used a 20-year time window moved from year to year starting in 1970 to maximize tree samples while having a sufficiently long period to test the effect of climate change. Of the 240 tree cores, we discarded three cores that were younger than 1970 and two other cores due to ring detection problems. For each time window and stand we built a network as described above and calculated its connectivity. Results with different window lengths provided qualitatively and quantitatively similar results. For each time window, we calculated the mean d.b.h. (i.e. mean tree size), the coefficient of variation of the d.b.h. (i.e. tree size heterogeneity) and the mean of the climatic variables described in *Climate data*.

#### Effect of land- and forest-use legacies and climate on tree growth synchrony

We analysed whether land- and forest use legacies, climate or both influenced tree growth synchrony from 1970 to 2020. For this aim, we fitted three linear mixed effects models of tree growth synchrony over 20-year time windows, assuming a normal distribution of errors and using stand identity as random effect. The first model included only climatic variables (i.e. total precipitation, number of heatwaves and number of late spring frosts) as additive fixed effects. The second model included only legacy type as a fixed effect, and the third model included both climate and legacy type as additive fixed effects.

To test the underlying drivers of land- and forest-use legacies modulating tree growth synchrony in response to climate change, we compared other three linear mixed models of tree growth synchrony over 20-year time windows also assuming a normal distribution of errors and using stand identity as random effect. One model including climatic variables and legacy type as additive fixed effects and a second model including climatic variables and forest structural diversity (i.e. mean tree size and tree size heterogeneity) as additive fixed effects. With these two models we tested whether legacy type or structural diversity was driving tree growth synchrony in response to climate change. We also included a third model considering climatic variables, legacy type and forest structural diversity as additive fixed effects to test whether there were other variables related to legacy type that were driving tree growth synchrony apart from structural diversity. All fixed predictors were standardised before being included in the models, the models were fitted using glmmTMB R package (Brooks et al., 2022) and compared in terms of AICc (Burnham & Anderson, 2002) using bbmle R package (Bolker & R Development Core Team, 2022). We also calculated goodness-of-fit using the pseudo-*R*^*2*^ described in Nakagawa & Schielzeth (2013) and implemented in MuMIn R package (Bartoń, 2022). The models’ fits were diagnosed using DHARMa R package (Hartig, 2022). All analyses were performed using R Statistical Software (v4.2.1; R Core Team 2022).

## RESULTS

### Land- and forest-use legacies and climate drive tree growth synchrony

Comparing the model containing land- and forest-use legacies and climatic variables (i.e. total precipitation, number of heatwaves and number of late spring frosts) with the model without climatic variables and the model without legacy type, the model containing both climatic variables and legacy type had the lowest AIC (*Δ*AICc legacy + climate = 0.0, *Δ*AICc climate = 15.5, *Δ*AICc legacy = 189.9). Moreover, climatic variables and legacy type explained similar and complementary portions of the tree growth synchrony variance (marginal *R*^*2*^ legacy + climate = 0.57, marginal *R*^*2*^ climate = 0.28, marginal *R*^*2*^ legacy = 0.29). Specifically, our results indicated more synchronous responses in dry years (Fig. 2A, C and Fig. 3), and with increases in the number of heatwaves and late spring frosts (Fig. 2A, D, E and Fig. 3). These results showed that both land- and forest-use legacies and climate determined tree growth synchrony in the period 1970-2020.

**Figure 2.**
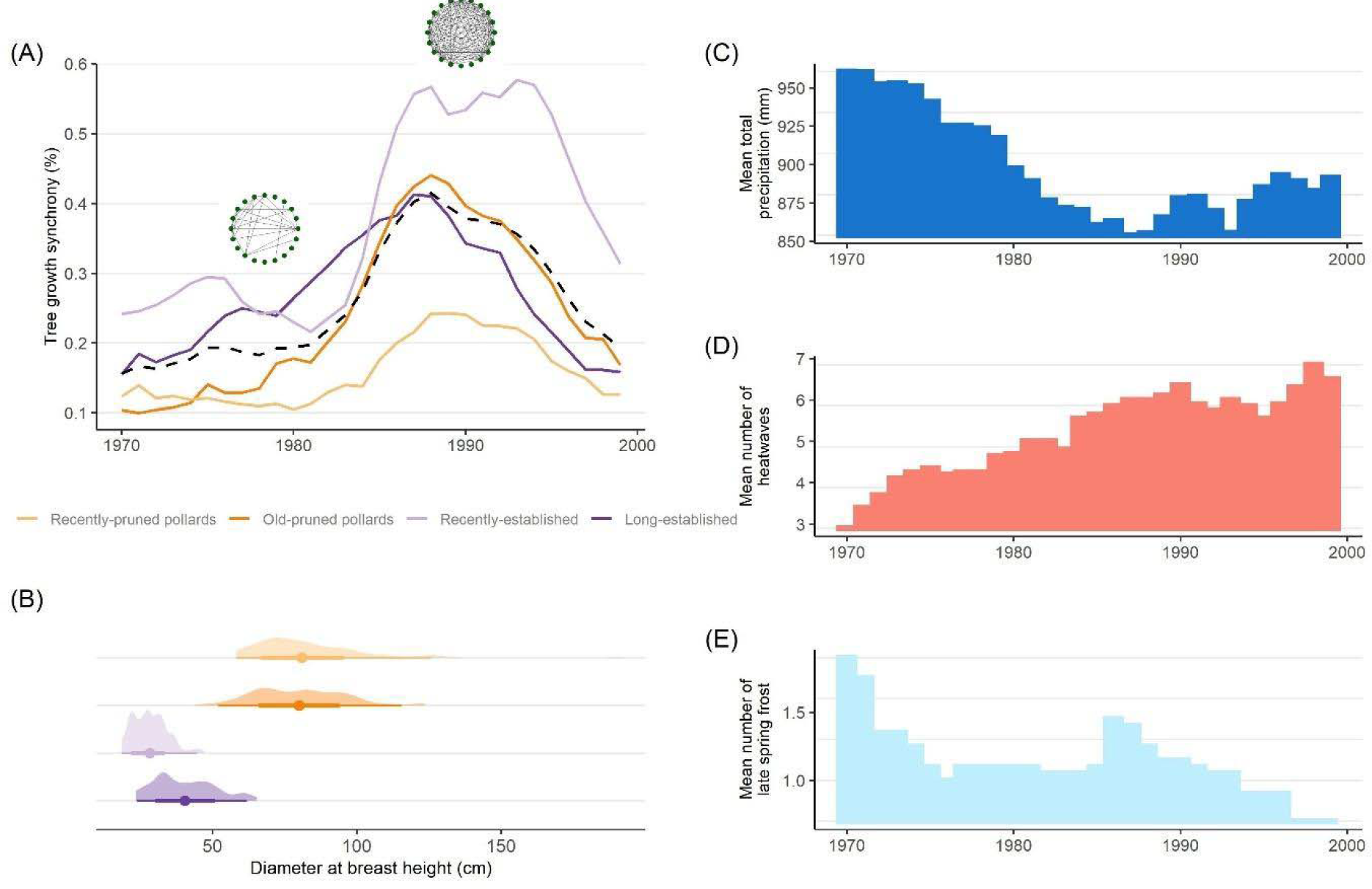
Tree growth synchrony as a function of land- and forest-use legacies, structural diversity of each legacy type and climate trends in the last decades. (A) Tree growth synchrony using 20-years moving windows between 1970-2020 as a function of land- and forest-used legacies. The solid lines represent the mean tree growth synchrony of each legacy type, and the dashed line represents the overall mean of tree growth synchrony of all legacy types. Two tree growth synchrony networks are shown, one in a low synchrony period (few links) and one in a high synchrony period (many links). Note that the two networks shown correspond only to one recently-established stand in two time windows and are used as an example to show the calculation and shape of tree growth synchrony networks. (B) Density and interval plot of the diameter at breast height of sampled trees where the point shows the mean tree size, and the 95% interval and density, the distribution of tree size of each legacy type. (C-E) Mean total precipitation, mean number of heatwaves and mean number of late spring frost using 20-years moving windows between 1970-2020.

**Figure 3.**
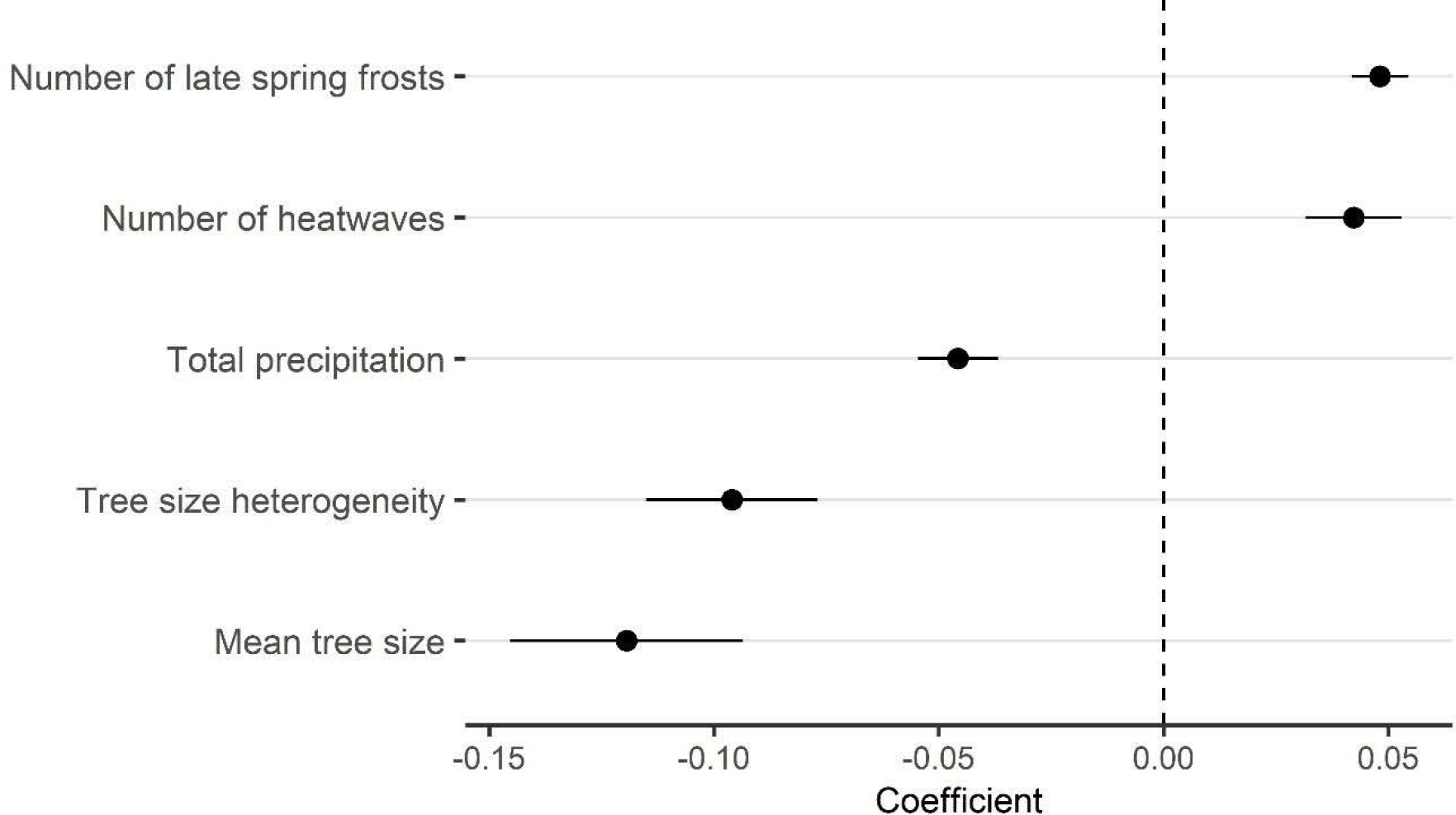
Sensitivity of tree growth synchrony to climate and forest structure. Estimated coefficients + 1 standard errors for the regression of tree growth synchrony on number of late spring frosts, number of heatwaves, total precipitation, tree size heterogeneity and mean tree size.

The magnitude of tree growth synchrony varied depending on legacy type, with the highest synchrony in recently-established stands and the lowest synchrony in recently-pruned pollards (Fig. 2). In addition, tree growth synchrony remained higher over time in recently-established stands than in other legacy types when precipitation increased and the number of heatwaves and late spring frosts decreased (see in Fig. 2 the decrease in tree growth synchrony *c*. 1990 for all legacy types except for recently-established stands).

### Forest structural diversity determines tree growth synchrony in response to climate

Comparing the model containing legacy type, climate and forest structural diversity with the model without legacy type and the model without forest structural diversity, we obtained that the lowest AIC was found in the full model and the model without legacy type (*Δ*AICc legacy + climate + structure = 0.0, *Δ*AICc climate + structure = 0.6, *Δ*AICc climate + legacy = 18.9). Since the difference *Δ*AICc was lower than two units, we selected the simpler model (i.e. climate + structure, marginal *R*^*2*^ = 0.53 and conditional *R*^*2*^ = 0.74). These results suggest that the underlying driver determining tree growth synchrony in response to climate is related to forest structural diversity (i.e. mean tree size and tree size heterogeneity). Our results showed more synchronous responses in dry years, with greater number of heatwaves and late spring frosts, and in stands with low tree size heterogeneity and mean tree size (Fig. 3).

## DISCUSSION

Our results showed an increase in tree growth synchrony in response to increasing climate change impacts in a widely distributed temperate tree species. Reduced total precipitation and increased temperature-related extreme events drove the observed increase in tree growth synchrony. However, land- and forest-use legacies modulated the magnitude of tree growth synchrony through changes in forest structural diversity (i.e. mean tree size and size heterogeneity). Our results, showing that recently-established forests had the greatest tree growth synchrony and that large trees and heterogeneous tree sizes decreased growth synchrony, have direct implications for forest management since heterogeneous structures could be favoured to adapt forests to novel climatic conditions.

We observed increases in tree growth synchrony in all sampled stands when precipitations decreased and the number of heatwaves and late spring frosts increased. This result confirms our first hypothesis and is consistent with previous studies that recorded increases in tree growth synchrony in response to increased climatic constraints for multiple species and forest types around the world (Boden et al., 2014; Gazol et al., 2020; Shestakova et al., 2016). Although climatic drivers of growth synchrony can vary depending on regional climate (Shestakova et al., 2016), the observed role of dry conditions and heatwaves agrees with the results obtained in populations of *F. sylvatica* in north-western Europe (Latte et al., 2015). Also, the effect of late frost frequency in tree synchrony is in line with the detrimental influence of late frosts in the performance of *F. sylvatica* in southern populations (Sangüesa-Barreda et al., 2021). Increased growth synchrony is related to high population vulnerability to climate change (Boden et al., 2014), which in the case of dominant species such as *F. sylvatica*, could jeopardise the stability and persistence of the entire ecosystem. Thus, our results contribute to previous observations pointing to increased vulnerability to climate change of temperate forests, especially those located at the southern distribution limits (Lindner et al., 2010; Millar & Stephenson, 2015).

The results obtained also support our second hypothesis, as stands with land- and forest-use legacies resulting in greater forest structural diversity showed less tree growth synchrony in response to climate change. Different drivers have been proposed to unravel why trees may respond differently to climate depending on past land- and forest-use, ranging from alterations in biomass (i.e. structural overshoot, Jump et al., 2017), in wood density (Alfaro-Sánchez et al., 2019) or in ectomycorrhizal communities (Correia et al., 2021; Mausolf et al., 2018). In this regard, our results suggest the key role of forest structural diversity in how anthropogenic legacies modulate forest vulnerabilities to climate change. The negative effect of tree size heterogeneity on tree growth synchrony indicates that structurally more heterogeneous forests may increase forest resilience by increasing individual-level growth variability in response to climate (Clark et al., 2012). In fact, different tree sizes may show different responses to climate (Day & Greenwood, 2011), spreading the risk of being negatively affected across different tree sizes.

The effect of mean tree size in growth suggests a high importance of ontogeny in assessing the impacts of climate change on forests (Heiland et al., 2022). Small trees –usually associated with young individuals– responded more synchronously to climate than large trees, highlighting the importance of maintaining old-growth forests to mitigate the negative effects of climate change on tree growth synchrony, which is a very relevant result in Europe, where old-growth forests are very scarce (Sabatini et al., 2018). Larger trees have larger carbohydrate reserves, more developed root system and greater connection to mycorrhizal networks than small trees. This could enhance tree growth resistance and resilience to climatic disturbances compared to younger and smaller trees (Ruiz-Benito et al., 2015; Zang et al., 2012). In addition, pollarded trees often occur at low densities, which can prevent strong inter-individual competition for resources that worsens the effects of climate change (Linares et al., 2010). Although our results showed that forest structural diversity explained a large part tree growth synchrony variance, other variables may also affect growth synchrony (Alfaro-Sánchez et al., 2019; Correia et al., 2021; Jump et al., 2017), suggesting that legacy effects might affect to different forest properties that can act synergistically in modulating the impact of climate change on tree growth.

We provide a novel method for estimating tree growth synchrony at individual level that opens a range of analysis possibilities previously unattainable. Our metric could be complemented with other more standard tree growth metrics to gain a comprehensive understanding of tree vulnerability to climate. Using this novel method, we found that recently-established stands had the highest tree growth synchrony. In addition, tree growth synchrony remained higher in recently-established stands over time than in other legacy types when precipitation began to increase and the number of heatwaves and late spring frosts started to decrease. These results suggest that, although recently-established forests are fundamental for carbon sink in the Iberian Peninsula (Vilà-Cabrera et al., 2017), they are vulnerable to climate change due to their more homogeneous structure leading to high tree growth synchrony. The high vulnerability of recently-established forests poses a risk to the carbon sink dynamics in Europe, which is mainly dominated by regrowing forests (Pugh et al., 2019) and where the first signs of carbon saturation are already visible (Nabuurs et al., 2013). Based on our results, recently-established forests could be handled by applying uneven-aged management options to increase forest structural diversity and decrease tree growth synchrony (Lafond et al., 2014). In addition, pollards (both recently- and old-pruned) showed lower synchrony than recently-established forests. Pruning reintroduction in old pollards often improves tree stability and vigour (Camarero et al., 2022), and is likely to be responsible for the lower growth synchrony observed in recently-pruned pollards compared to old-pruned pollards. Recovering tree pollarding and associated wood pastures could bring additional benefits beyond reducing tree growth synchrony, such as, reduced inter-individual tree competition, high biodiversity characteristic of these ecosystems and compatibility of livestock activities and forest-use (Sjölund & Jump, 2013). Our results therefore highlight the most climate-vulnerable forests based on past land- and forest-use, but also show key indications for management actions to adapt forests to the increasing impacts of climate change.

## ACKNOWLEDGEMENTS

This study was funded by the municipality of Oñati (Basque Country, Spain). PRB and JTT were supported by the Community of Madrid Region under the framework of the multi-year Agreement with the University of Alcalá (Stimulus to Excellence for Permanent University Professors, EPU-INV/2020/010). JA, PRB, JTT, MAZ and AH are funded by the Science and Innovation Ministry (subproject LARGE, Nº PID2021-123675OB-C41). EA is supported by the grant PID2019-110470RA-100 (ADAPTAMIX) funded by MCIN/AEI/10.13039/501100011033. AH was supported by the Basque Country Government funding support to FisioClima CO_2_ (IT1682-22) research group. We thank F. Rodríguez-Sánchez, V. Cruz-Alonso, J. M. Serra-Diaz and P. Lloret for the interesting discussion on the manuscript and their useful comments to improve it.

